# Stable isotopic composition of Antarctic and Patagonian marine mammals collected before and during industrial-scale whaling: assessing the baseline for long-term changes in the marine ecosystem

**DOI:** 10.1101/2024.08.13.607746

**Authors:** Evgeny Genelt-Yanovskiy, Anna Genelt-Yanovskaya, Maria Fontanals-Coll, Kweku Afrifa Yamoah, Oliver Craig, Richard Sabin, James Scourse

**Affiliations:** Department of Earth and Environmental Science, University of Exeter, Penryn Campus, Cornwall, TR10 9FE UK; University of York, BioArCh, Environment Building 2nd Floor, Wentworth Way, Heslington, York, YO10 5DD, UK; The Natural History Museum, Cromwell Road, London SW7 5BD, UK

**Keywords:** Antarctic, cetacean, pinniped, bone collagen, museum collections, carbon, nitrogen, stable isotopes, Suess effect, trophic position, foraging ecology

## Abstract

Great Antarctic expeditions, seal hunting and whaling industries left a legacy in natural history collections. To provide the basis for analysing the impact of whaling on marine ecosystem structuring, we conducted the bulk isotope analysis from the specimens of baleen whales (*Balaenoptera musculus* and *B. physalus*), and seals *(Arctocephalus australis* and *Hydrurga leptonyx*) collected between 1843 to 1951 from the South Atlantic, Patagonian waters, Southern Ocean and Antarctic coastal seas, and preserved in the collection of Natural History Museum, London. Analysis of this material indicates the pre-industrial whaling state of these environments, and changes in the trophic position of whales and seals during the period of extensive human pressure. Having controlled for the Suess effect, δ^13^C values in *B. musculus*, *B. physalus* and *H. leptonyx* were different before and after the onset of industrial-scale whaling (1904). Bone collagen δ^15^N values and corresponding trophic position indicate possible trophic changes in *A. australis,* and variability of the foraging areas of *B. musculus*. This study highlights the use of museum specimens for tracing historical trends associated with changes in the population structure and distribution of species and which indicate long-term variability in their foraging ecology.

## Introduction

The massive numbers of marine mammals and seabirds is among the most notable components of the Antarctic ecosystem. They play an important role as highly mobile consumers connecting the open sea pelagic and nearshore marine food webs. Antarctic and Subantarctic waters are crucial feeding grounds for many cetaceans, including baleen whales, attracted by seasonally high densities of krill (1,2). Mainland Antarctica and the Subantarctic islands houses vast breeding colonies of pinnipeds, and the productivity of the region is reflected in estimates that it contains more than a half of the global seal population.

Mammals function as keystone species, and serious depletion of their numbers can cause major changes in ecosystem functioning. Historically, overkill was the most important human activity affecting the abundance of marine mammals. Antarctic seas were effectively pristine prior to discovery of the continent in the eighteenth century. Whaling and sealing spread alongside the exploration of the region. Throughout the nineteenth and early twentieth centuries, the Antarctic was viewed as an almost limitless source of marine mammals to be hunted for skins, oil, and other products. The development of seal hunting and subsequently whaling resulted in a severe decline in commercially important species. Seal hunting brought the Antarctic fur seal *Arctocephalus gazella* almost to extinction by the late 1700-early 1800s in South Georgia, and the whaling industry peaked in the early 1900s when the permanent whaling stations were opened at Grytviken (South Georgia) and Deception Island (South Shetland Islands) (3,4). Following international moratoria on industrial whaling activities, whale populations have been recovering over recent decades after a long period of decline, but recent census studies indicate that their abundance has still not attained pre-whaling numbers (5,6).

The significant decline in the abundance of marine mammals in the late 19^th^ – early 20^th^ centuries caused by industrial-scale exploitation may well have had a significant impact on the marine ecosystem of Antarctica, as well as on the structure and distribution of populations of the species affected. Intensive whaling and seal hunting in Antarctica coincided with the period known as “Heroic Age of Antarctic exploration”, a series of naval and land expeditions between the late 19th century and 1920s. This period left a legacy in natural history museum collections around the world, including the specimens of species targeted by industrial whaling and sealing. Museum specimens are increasingly important for gaining ecological and biological data, and as carriers of geochemical and molecular information, important for studying rare, elusive and even extinct species. Historical museum “specimens of opportunity” are often challenging to use due to uncertainties in time and location of collections, and in the quality of post-collection curation, but similarly to the subfossil specimens from excavations of archaeological or paleontological sites can provide snapshots of past genetic diversity, spatial origin and diet (7–9). Stable isotope analysis, most commonly the bulk δ^13^C and δ^15^N isotopes of marine species, can indicate latitudinal range shifts, migrations between nearshore and offshore environments as well as changes in trophic structure (10).

Historical specimens of Antarctic whales from various collections have become increasingly important in isotopic and genetic studies, being used to describe niche partitioning between fin (*Balaenoptera physalus)* and sei (*B. borealis*) whales (11), changes in mitochondrial DNA diversity in blue (*B. musculus)*, humpback (*Megaptera novaeangliae*) and fin whales (12). Isotopic analyses of collections representing Antarctic fur seal (*A. gazella*) in Antarctica (13) and South American fur seal (*A. australis*) in Patagonia (14) indicate long-term latitudinal and longitudinal shifts in their prey distributions. The stable isotopic analysis approach is based on the relationship that carbon and nitrogen isotopes in the predator tissues reflect those of its prey in a predictable way (15). Factors such as nutrient resources, composition of primary producers and regional geographic characteristics result in distinct isotopic patterns between different regions (16), allowing the use of stable isotope analysis for investigating large-scale movements of predators (16,17).

In this study, we measured δ^13^C and δ^15^N compositions of the oldest Antarctic and Subantarctic marine mammal bone specimens of opportunity preserved in the Natural History Museum in London (NHMUK). These specimens represent the species most targeted by sealing and whaling activity and thus collected during the expeditions prior to, and during, the development of industrial scale harvesting. Following the timeline and southward trend of exploration of Subantarctic and Antarctic waters, the oldest specimens in the dataset correspond to the period of survey of the Strait of Magellan by the HMS *Alert* Expedition (1876–1884), and, the James Clark Ross Expedition onboard HMS *Erebus* and HMS *Terror* (1838–1843). By focusing primarily on baleen whales - blue whale *B. musculus* and fin whale *B. physalus*, and two species of seals – leopard seal *Hydrurga leptonyx* and South American fur seal *A. australis* - we explore the following research questions: (i) How have δ^13^C and δ^15^N stable isotopes varied during the onset of intensive whaling; (ii) Can the observed variability in δ^13^C and δ^15^N be used for fingerprinting of foraging areas in highly mobile marine mammals, (iii) Can the observed trends in δ^15^N indicate spatial and temporal changes in the trophic position of the species.

## Materials and methods

### Sampling

Samples were collected at the mammal collection of Natural History Museum, London (NHMUK) The two key criteria for the selection of samples from the NHMUK collection were date of sampling and geographical origin identified either from the museum label directly or based on records related to the research cruise or programme. While the main focus of the study was on the South Georgia - Subantarctic islands biogeographical province according to e.g. (18) and waters surrounding Antarctic Peninsula (Maritime Antarctic), whale and seal specimens from Falkland Islands and Patagonia (Southern cool temperate province) were also considered, as these areas were visited during the Antarctic Expeditions and are, to a great extent, influenced by wider Antarctic climate system (19,20). In this study, we relied on species identification by the NHMUK, which included the DNA barcoding of all the fin and blue whale specimens implemented at NHMUK facility. Before sampling, the surface of the target fragment of the bone was cleaned using the Dremel tool with the bristle polishing brush bit, and the cutting of bone fragment was implemented using handheld cutting tools and a Dremel with either dental bur or carbide cutting disk. In total, 36 samples of bone fragments were analysed (Table 1).

**Table 1.**
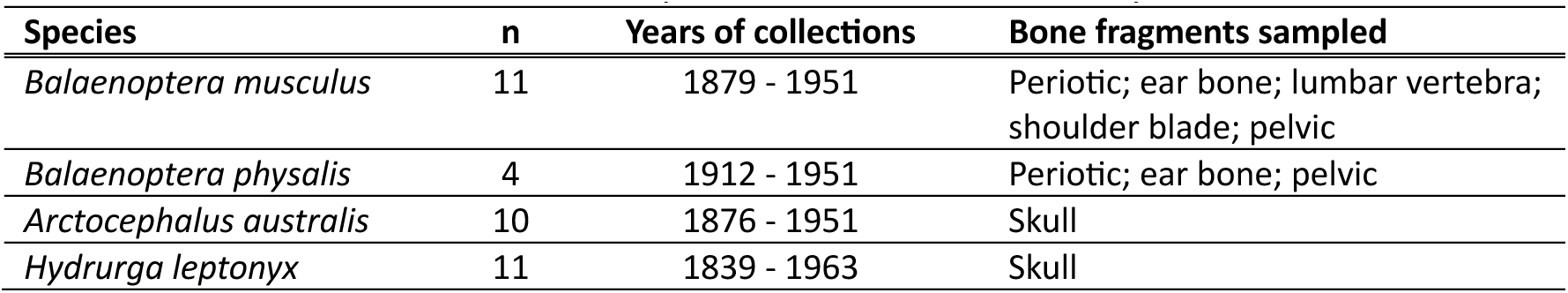
Bone material and number of specimens used in the study.

The initial weight of the bone samples was about 500 mg. Collagen was extracted and prepared for stable isotope analysis at the BioArCh laboratories, University of York (UK). The bone samples (approx. 200 to 300 mg) were demineralized using 0.4 M HCl at 4°C for several weeks, then rinsed with deionised water (milli-Q) and gelatinised with 0.001 M HCl at 70°C for 24 to 48 h. The supernatant containing the collagen was filtered using polyethylene Ezee filters (Elkay Laboratories, 9 ml, pore size 60–90 µm) and then 30 kDa Amicon Ultra-4 centrifugal filter units (Millipore, MA, USA). Samples were then frozen for 24–48 h at −20°C, lyophilised, and weighed into tin capsules (0.4-0.6 mg per duplicate) for stable isotope analysis.

### Bulk stable isotope analysis

The BioArCh laboratories at the University of York’s Department of Archaeology conducted duplicate measurements of stable carbon and nitrogen isotope ratios. Using a Sercon EA-GSL elemental analyzer linked to a Sercon 20-22 continuous flow isotope ratio mass spectrometer (Sercon, Crewe, UK), the ratios of ^13^C:^12^C and ^15^N:^14^N were assessed relative to standards (V- PDB for carbon and AIR for nitrogen) and reported in delta notation (δ) in parts per mil (‰). Stable carbon and nitrogen isotope ratios were measured using a Sercon HS2022 continuous flow isotope ratio mass spectrometer at BioArCh, University of York (UK).

Stable carbon and nitrogen isotope ratios were calibrated relative to VPDB and AIR scales using the standard reference materials IsoAnalytical Cane (δ^13^C = -11.64‰ ± 0.03), Sigma Methionine (δ^13^C = -35.83‰ ± 0.03, δ^15^N = -0.76‰ ± 0.05) and IAEA N2 (δ^15^N = 20.41‰ ± 0.12). Uncertainty was monitored using the standard reference materials Sigma fish gel (δ13C = -15.27‰ ± 0.04, δ15N = 15.21‰ ± 0.13), IsoAnalytical Alanine (δ^13^C = -23.33‰ ± 0.10, δ^15^N = -5.56‰ ± 0.14) and IsoAnalytical Soy (δ^13^C = -25.22‰ ± 0.03, δ^15^N = 0.99‰ ± 0.07). Precision (u(Rw)) was determined to be ± 0.165‰ for δ^13^C and ± 0.124‰ for δ^15^N based on repeated measurement of calibration standards, check standards and sample replicates. According to calibration and check standards (Table 1 Supplementary Material 1), measurement precision (ssrm, pooled measured standard deviation of calibration and check standards) was ± 0.151‰ for δ^13^C (df = 28) and ± 0.124‰ for δ^15^N (df = 24). All the samples (42/42) were analysed in duplicate (Table 2 Supplementary Material 1). The measurement precision specific to samples (srep, pooled standard deviation of sample replicates) was ± 0.094‰ for δ^13^C (df = 47) and ± 0.050‰ for δ^15^N (df = 47). Accuracy (u(bias)) was determined to be ± 0.091‰ for δ^13^C and ± 0.159‰ for δ^15^N based on the difference between observed and reported δ values (RMSbias) and long-term standard deviations (u(Cref)) of check standards. Total analytical standard uncertainty (uc) was estimated to be ± 0.188‰ for δ^13^C and ± 0.204‰ for δ^15^N (21).

**Table 2.** Oceanic Suess Effect correction values extracted from doi:10.1594/PANGAEA.872004 (see Eide et al., 2017 for details). N – number of point estimates of total accumulated Suess Effect in surface (0 meter) water layer. Total CSE – mean value of total oceanic Suess Effect for each region. The a-value – expected per-year δ^13^C decline (‰).

|  | Latitude range |  | Longitude range |  | n | Total CSE | a-value |
| --- | --- | --- | --- | --- | --- | --- | --- |
| Patagonia / Strait of Magellan | -49 | -55 | -77 | -67 | 19 | -0.91 | -0.0065 |
| Falkland Islands | -50.8 | -52.5 | -62 | -57.4 | 2 | -0.93 | -0.0066 |
| South Georgia | -53 | -56 | -38.7 | -33.9 | 13 | -0.73 | -0.0052 |
| South Shetlands | -61 | -63 | -63 | -57 | 9 | -0.75 | -0.0054 |
| Ross Sea | -71 | -77 | 171 | -160 | 173 | -0.71 | -0.0051 |

### Data analysis

Basic descriptive statistics to explore variation for both δ^13^C and δ^15^N within the bone samples were performed using R (R Core Team, 2024) with RStudio (RStudio Team, 2024). Raw δ^13^C values were normalized for potential lipid contamination. The resulting δ^13^C_normalized_ provides an estimate of δ^13^C that is normalized for the effects of lipid concentration on δ^13^C and is comparable to the δ^13^C after direct chemical lipid extraction (22).

Obtained δ^13^C_normalized_ values were then corrected for the oceanic Suess Effect – the depletion in the δ^13^C isotopic composition of dissolved inorganic carbon (DIC) in oceans due to the increase in anthropogenic CO_2_ released into the atmosphere since the Industrial Revolution (Keeling, 1979; Misarti et al., 2009; Keeling et al., 2017). Calculations were performed using the following equation introduced by Hilton et al. (2006), modified by Misarti et al. (2009) and later presented within the SuessR package (23): Suess Effect correction factor = ***a*** * exp(***b***\*0.027)

Where ***a*** is a constant reflecting the maximum observed rate of δ^13^C decline in surface waters DIC in a specific region, ***b*** is the year of sample collection minus 1850 (taken as the onset of the Industrial Revolution) and 0.027 is the parameter value obtained by Hilton et al. (2006) after fitting an exponential curve to the global ocean δ^13^C data from 1945 to 1997 published by Gruber et al. (1999). The values of total accumulation of Suess Effect for the constant ***a*** were obtained from Eide et al. (2017) for the surface waters (0 meters depth layer in a database available under the doi:10.1594/PANGAEA.872004). These values were obtained and averaged for five regions (Table 2). Per-year ratios of ***a*** were obtained by dividing these values by 140, i.e., the number of years between the year of 1850 (Industrial Revolution) and 1990 (start of the main decade of collecting data in (24)).

Bone δ^15^N signatures were converted to trophic position (TP) using the following equations (25,26):

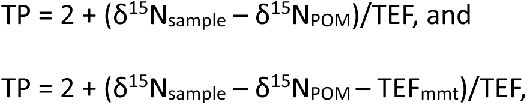

where δ^15^N_sample_ is nitrogen value in marine mammal bone, POM is δ^15^N value of marine particulate organic matter, and TEF/TDF is trophic enrichment (determination) factor (2 – 3.4) in δ^15^N for marine mammals (25,27–31). TEF can be highly variable among species and foraging areas, and thus is difficult to estimate in large marine mammals who migrate long distances. We used two approaches for calculating TEF for baleen whales following recent publications (28). The first uses TEF values previously obtained for the fin whale (27), later called as ‘Borrell TEF’. The second is typically accepted and widely used for marine ecosystem TEF values from (31), hereafter ‘Post TEF’. In the equation above, TEF_mmt_ is the tissue-specific trophic enrichment factor in δ^15^N, for baleen whale bone protein samples TEF_mmt_ = 2.03 ± 0.71 in was used (32).

Since the isotopic signatures of POM vary spatially and temporally in the study region (33), observed ranges in POM isotopic signature corresponding to the location of the specimen find site were used. Data on POM were obtained from previously published modern datasets; δ^15^N_POM_ = 0.6 for Ross Sea, δ^15^N_POM_ = 1.7 for South Shetlands, δ^15^N_POM_ = 1.5 for the South Atlantic sector of the Southern Ocean, δ^15^N_POM_ = 3.6 for South Georgia (range 2 – 5.4), δ^15^N_POM_ = 8.1 for Strait of Magellan (range 5.6 – 12.2), δ^15^N_POM_ = 9.4 for Falkland Islands, δ^15^N_POM_ = 9.4 for Patagonian West Coast (34–39).

Consumer and source isotope data were analysed using the MixSIAR package in R (40–42), where source data were composed of mean and standard deviation (SD) from potential prey sources. To build the potential regional dietary database for the blue whale (Table 3), stable isotopic signatures of dominant krill species were obtained for five regions, from Antarctic shelf waters to Patagonian shelf waters (43–47). Isotopic Data on potential prey objects of South American fur seal (Table 3) were obtained based on recent studies of behavior and dietary preferences of the species, and isotopic signatures were then mined from literature (43,48,49). All the used δ^13^C values of potential prey items passed the same Suess-correction procedures as the marine mammal data, the difference between non-corrected and corrected values for the dietary source data varied between 0.29 and 0.38‰.

**Table 3.**
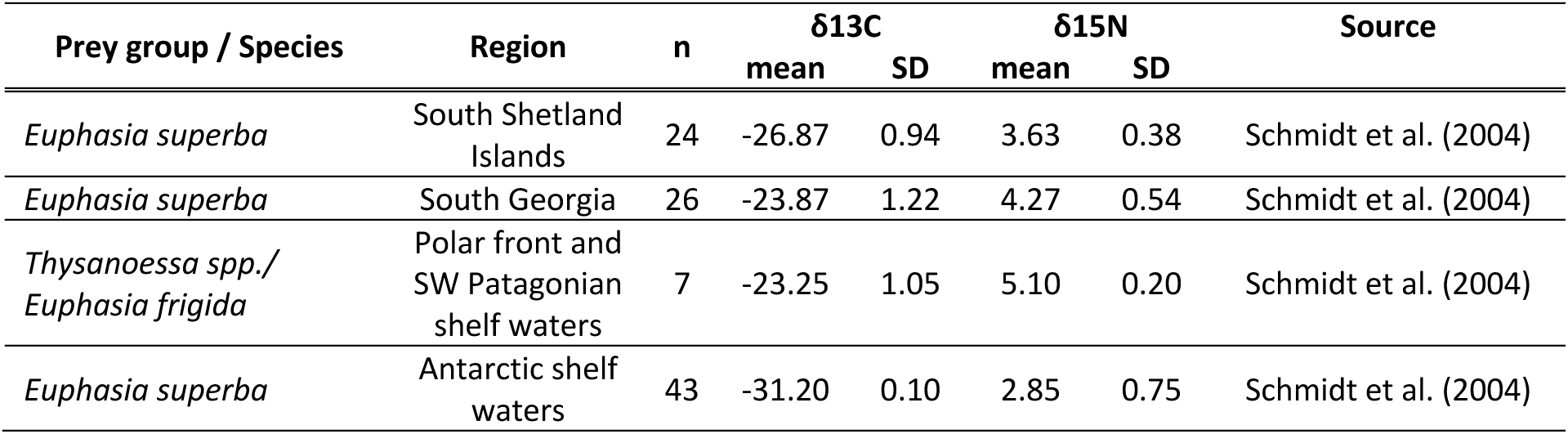
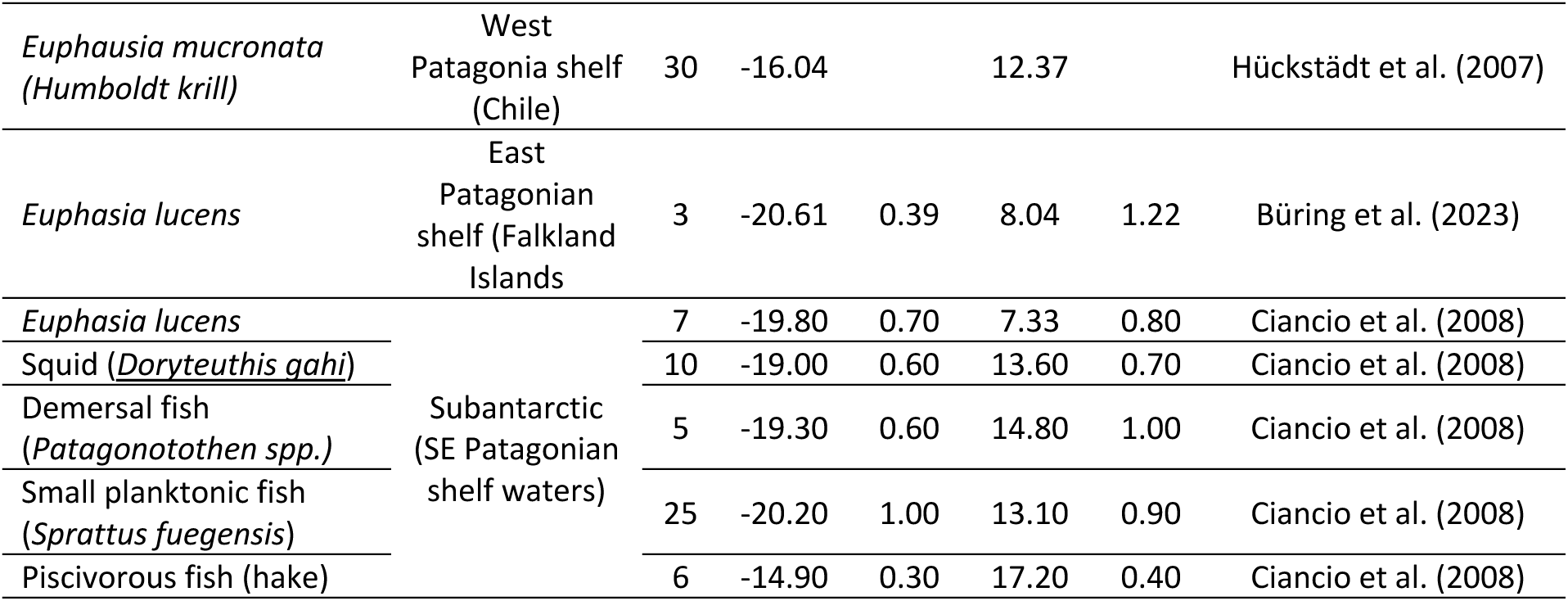
Stable isotope signatures of potential prey items for the blue whale and South American fur seal, mined from the literature (see text for references) and used for the MixSIAR modeling.

The models ran three Markov Chain Monte Carlo (MCMC) chains with 100000 iterations, a burn-in of 50000, and were thinned by 50. MixSIAR reports results as mean, SD and credible intervals for each posterior density distribution per prey source. To identify the within-group variability in dietary sources, SIMPER analysis was implemented in PAST v5.0 (50) using the matrix of obtained mean values of modelled proportion of diet. PAST v5.0 was also used to run PERMANOVA analysis applied to various among-group comparisons.

## Results

### Stable isotope data

The carbon to nitrogen (C:N) atomic ratio range from 3.29 to 4.3 in the analysed marine mammal specimens. Several specimens demonstrated signatures of lipid contamination based on the C:N mass values (Table 1 Supplementary Material 2). Thus, a mathematical correction (see the Materials and Methods for details) was applied to raw δ^13^C values for all specimens before subsequent analyses. The δ^13^C_normalised_ and δ^15^N values vary significantly among both blue whale and fin whale historical specimens studied (Figure 2, Table 4). The δ^13^C_normalised_ values of blue whale specimens vary between -22.76‰ to -13.14‰, and similarly δ^15^N values vary between 5.26‰ and 17.05‰. In fin whale, δ^13^C_normalised_ values range between -21.17‰ to -15.27‰, and δ^15^N values range between 6.13‰ to 11.64‰. The δ^13^C_normalised_ values in South American fur seals demonstrate a narrow range between -12.99‰ to -11.49‰, while δ^15^N values vary between 16.73‰ to 23.56‰. In leopard seals, δ^13^C_normalised_ values showed more pronounced variation (ࢤ22.01‰ to -15.83‰), compared to variation in δ^15^N between 7.45‰ and 11.27‰.

**Figure 1.**
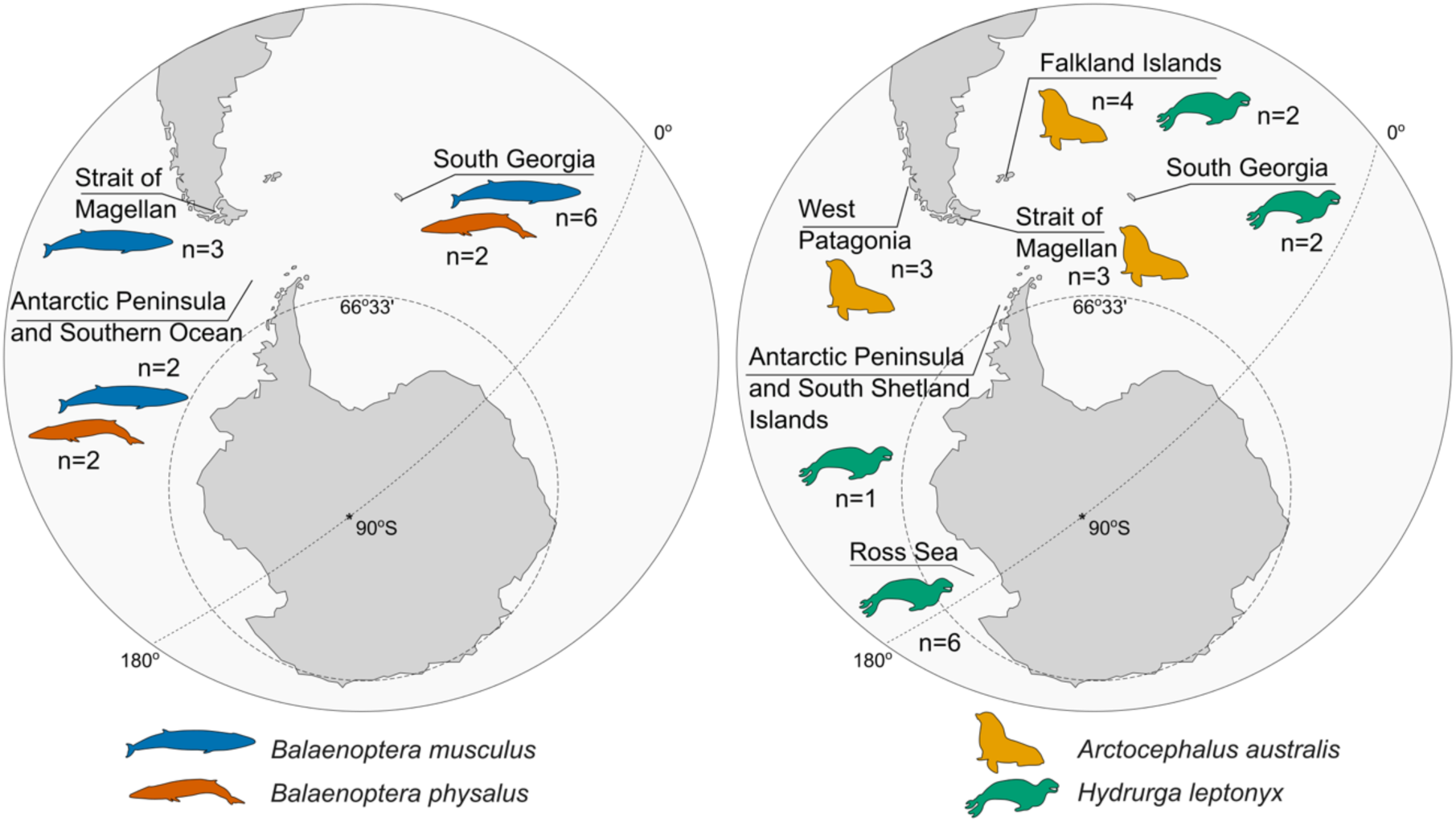
Locations of samples of marine mammals analysed in the study. Map redrawn from Google Earth. Detailed description of locations and NHM accession numbers are available in Table 1 Supplementary Material 2.

**Figure 2.**
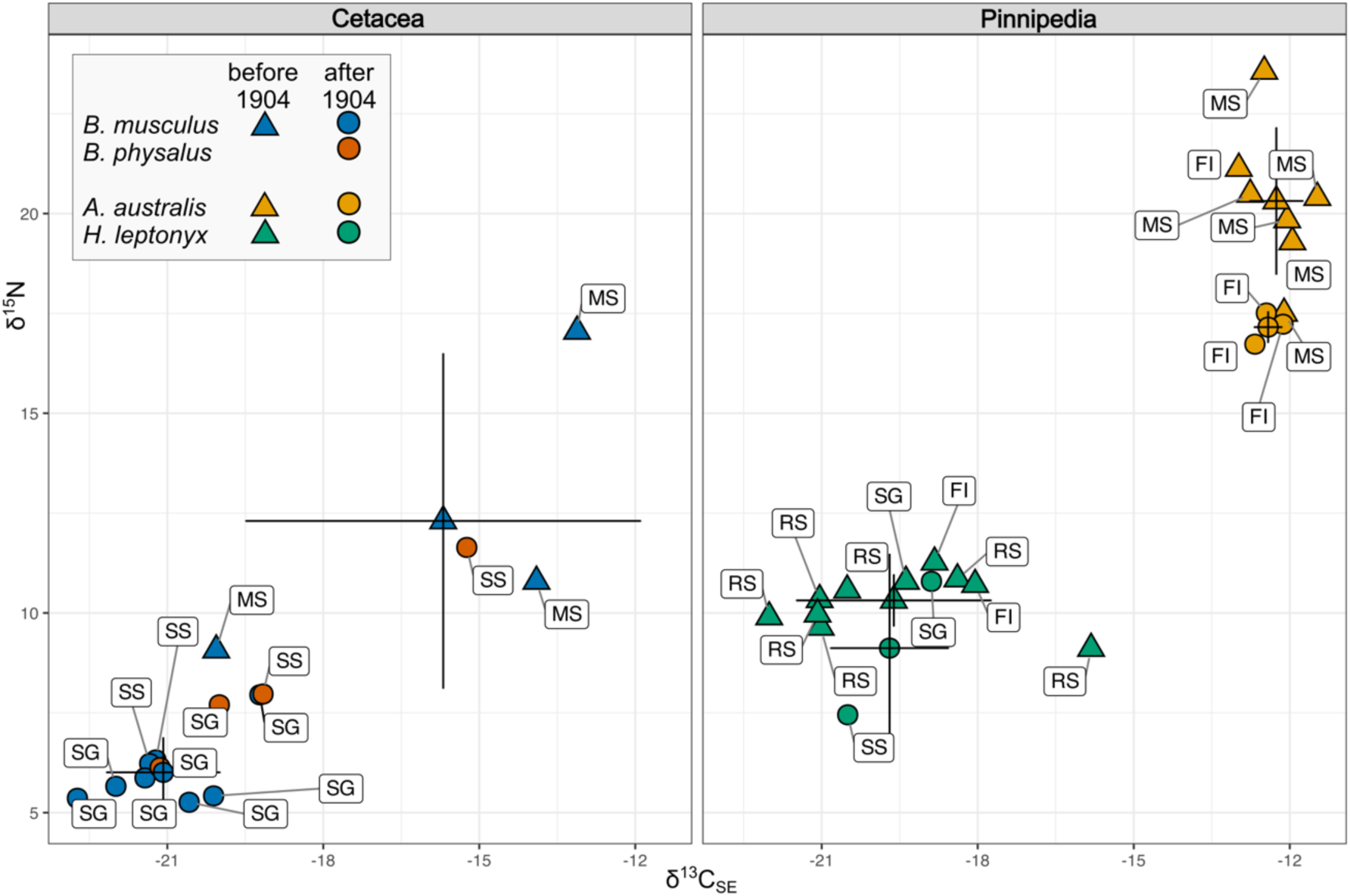
Suess corrected δ^13^C and δ^15^N biplot summarizing within-species variability of samples collected before (triangles) and after (circles) the opening of whaling station at South Georgia in 1904. Points with error bars denote mean±SD values for each period. Region codes: MS – Strait of Magellan and Patagonian West Coast, FI – Falkland Islands, SG – South Georgia, SS – Southern Ocean and South Shetland Islands, RS – Ross Sea.

**Table 4.** Summary of stable isotope values (Mean±SD) of bone collagen samples of marine mammals collected between the 1880s and 1960s.

| species | n | Raw $\delta^{13}\text{C} \pm \text{SD}$ | $\delta^{13}\text{C}_{\text{normalised}} \pm \text{SD}$ | $\delta^{13}\text{C}_{\text{SE}} \pm \text{SD}$ | $\delta^{15}\text{N} \pm \text{SD}$ |
| --- | --- | --- | --- | --- | --- |
| *Balaenoptera musculus*<br>blue whale | 11 | -19.86 ± 3.32 | -19.64 ± 3.18 | -19.61 ± 3.17 | 7.73 ± 3.57 |
| *Balaenoptera physalus*<br>fin whale | 4 | -19.10 ± 2.64 | -18.91 ± 2.56 | -18.88 ± 2.26 | 8.36 ± 2.33 |
| *Arctocephalus australis*<br>South American fur seal | 10 | -12.31 ± 0.48 | -12.33 ± 0.45 | -12.31 ± 0.45 | 19.37 ± 1.25 |
| *Hydrurga leptonyx*<br>leopard seal | 11 | -20.09 ± 1.25 | -20.00 ± 1.30 | - 19.97 ± 1.31 | 10.21 ± 1.04 |

The Suess-corrected δ^13^C values demonstrate the same trends and do not deviate more than 0.11‰ from δ^13^C_normalised_ values, and the mean value for all species is 0.027‰ (Figure 2, Table 4). The maximum correction value of the raw δ^13^C to take into account the potential effects of lipid contamination and oceanic Suess Effect (0.97‰) was applied to the blue whale specimen from South Georgia collected in 1914 (ZC.1953.12.1.16). The sample from this specimen is characterized by a very high C:N mass ratio (4.3), and thus the raw δ^13^C value 23.7‰ has been corrected to -22.73‰. The complete set of individual δ^13^C and δ^15^N values, spatial variation of indices and detailed information about the origin of each specimen analysed in this study is presented in Table 1 Supplementary Material 2.

### pre- and post-1904 stable isotopic values

In total, 19 specimens pre-dating the opening of the whaling station at South Georgia were analyzed, including three specimens of blue whale, seven specimens of South American fur seal, and nine specimens of leopard seal (Table 5, Figure 2). The mean (± SD) Suess corrected δ^13^CSE value over this period in blue whale samples is -15.69 ± 3.8, and mean δ^15^N = 12.3 ± 4.2, in South American fur seal samples mean δ^13^C_SE_ = -12.26 ± 0.52 and mean δ^15^N = 20.32 ± 1.84, and leopard seal mean δ^13^C_SE_ = -20.04 ± 1.40 and mean δ^15^N = 10.39 ± 0.65 (Table 5).

**Table 5.**
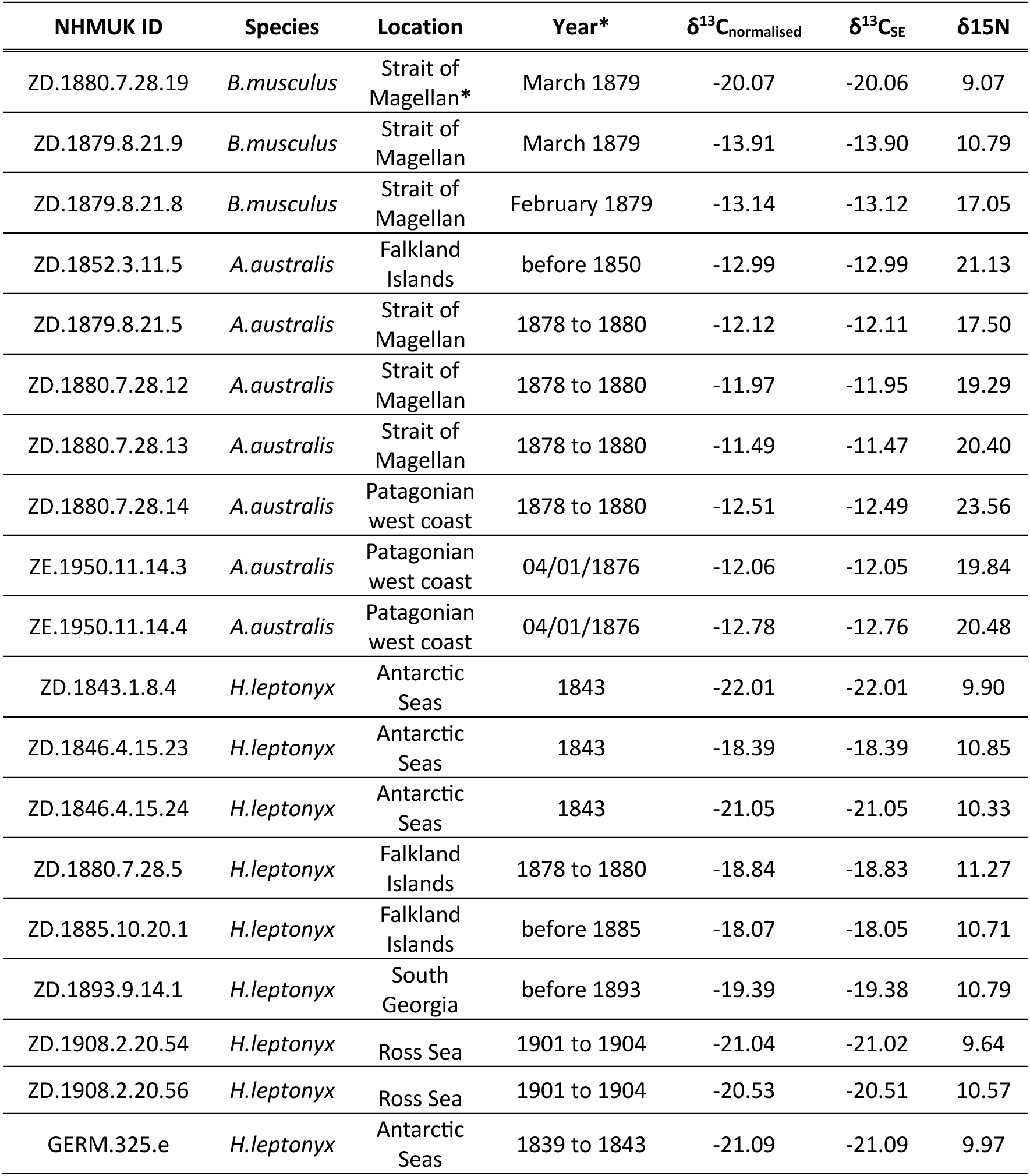
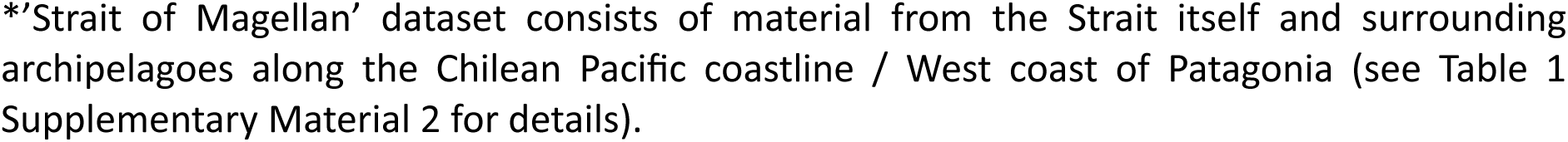
Stable isotope ratios in bone collagen of NHM specimens pre-dating the opening of the whaling station at South Georgia in 1904.

| NHMUK ID | Species | Location | Year* | $\delta^{13}\text{C}_{\text{normalised}}$ | $\delta^{13}\text{C}_{\text{SE}}$ | $\delta^{15}\text{N}$ |
| --- | --- | --- | --- | --- | --- | --- |
| ZD.1880.7.28.19 | *B.musculus* | Strait of Magellan* | March 1879 | -20.07 | -20.06 | 9.07 |
| ZD.1879.8.21.9 | *B.musculus* | Strait of Magellan | March 1879 | -13.91 | -13.90 | 10.79 |
| ZD.1879.8.21.8 | *B.musculus* | Strait of Magellan | February 1879 | -13.14 | -13.12 | 17.05 |
| ZD.1852.3.11.5 | *A.australis* | Falkland Islands | before 1850 | -12.99 | -12.99 | 21.13 |
| ZD.1879.8.21.5 | *A.australis* | Strait of Magellan | 1878 to 1880 | -12.12 | -12.11 | 17.50 |
| ZD.1880.7.28.12 | *A.australis* | Strait of Magellan | 1878 to 1880 | -11.97 | -11.95 | 19.29 |
| ZD.1880.7.28.13 | *A.australis* | Strait of Magellan | 1878 to 1880 | -11.49 | -11.47 | 20.40 |
| ZD.1880.7.28.14 | *A.australis* | Patagonian west coast | 1878 to 1880 | -12.51 | -12.49 | 23.56 |
| ZE.1950.11.14.3 | *A.australis* | Patagonian west coast | 04/01/1876 | -12.06 | -12.05 | 19.84 |
| ZE.1950.11.14.4 | *A.australis* | Patagonian west coast | 04/01/1876 | -12.78 | -12.76 | 20.48 |
| ZD.1843.1.8.4 | *H.leptonyx* | Antarctic Seas | 1843 | -22.01 | -22.01 | 9.90 |
| ZD.1846.4.15.23 | *H.leptonyx* | Antarctic Seas | 1843 | -18.39 | -18.39 | 10.85 |
| ZD.1846.4.15.24 | *H.leptonyx* | Antarctic Seas | 1843 | -21.05 | -21.05 | 10.33 |
| ZD.1880.7.28.5 | *H.leptonyx* | Falkland Islands | 1878 to 1880 | -18.84 | -18.83 | 11.27 |
| ZD.1885.10.20.1 | *H.leptonyx* | Falkland Islands | before 1885 | -18.07 | -18.05 | 10.71 |
| ZD.1893.9.14.1 | *H.leptonyx* | South Georgia | before 1893 | -19.39 | -19.38 | 10.79 |
| ZD.1908.2.20.54 | *H.leptonyx* | Ross Sea | 1901 to 1904 | -21.04 | -21.02 | 9.64 |
| ZD.1908.2.20.56 | *H.leptonyx* | Ross Sea | 1901 to 1904 | -20.53 | -20.51 | 10.57 |
| GERM.325.e | *H.leptonyx* | Antarctic Seas | 1839 to 1843 | -21.09 | -21.09 | 9.97 |

\*'Strait of Magellan' dataset consists of material from the Strait itself and surrounding archipelagoes along the Chilean Pacific coastline / West coast of Patagonia (see Table 1 Supplementary Material 2 for details).

Among the post-1904 specimens (Figure 2), the δ^13^C_SE_ mean value for blue whale is -21.08 ± 1.1, and the mean δ^15^N value 6.01 ± 0.88 (N = 8). In South American fur seals, the mean δ^13^C_SE_ value is -12.26 ± 0.52 and δ^15^N mean value 17.16 ± 0.40 (N=3). In leopard seals, the mean δ^13^C_SE_ value is 19.70 ± 1.14 and δ^15^N 9.29 ± 1.70 (N = 2).

Differences were most pronounced in the δ^15^N values for South American fur seal (PERMANOVA F=14.61, p = 0.041) (Figure 3), yet no difference in δ^13^C_SE_ values for this species has been found between the samples collected in the end of the 19^th^ and in early 20^th^ centuries. Despite the large variability in δ^13^C_SE_ values in general, no significant changes in either δ^13^C_SE_ or δ^15^N in leopard seals have been found. Changes in stable isotope signatures after 1904 in blue whale are significant for both carbon and nitrogen with more pronounced changes in δ^15^N (PERMANOVA F=38.94, p=0.0011); differences in δ^13^C_SE_ are also significant (PERMANOVA F=15.02, p=0.0134).

**Figure 3.**
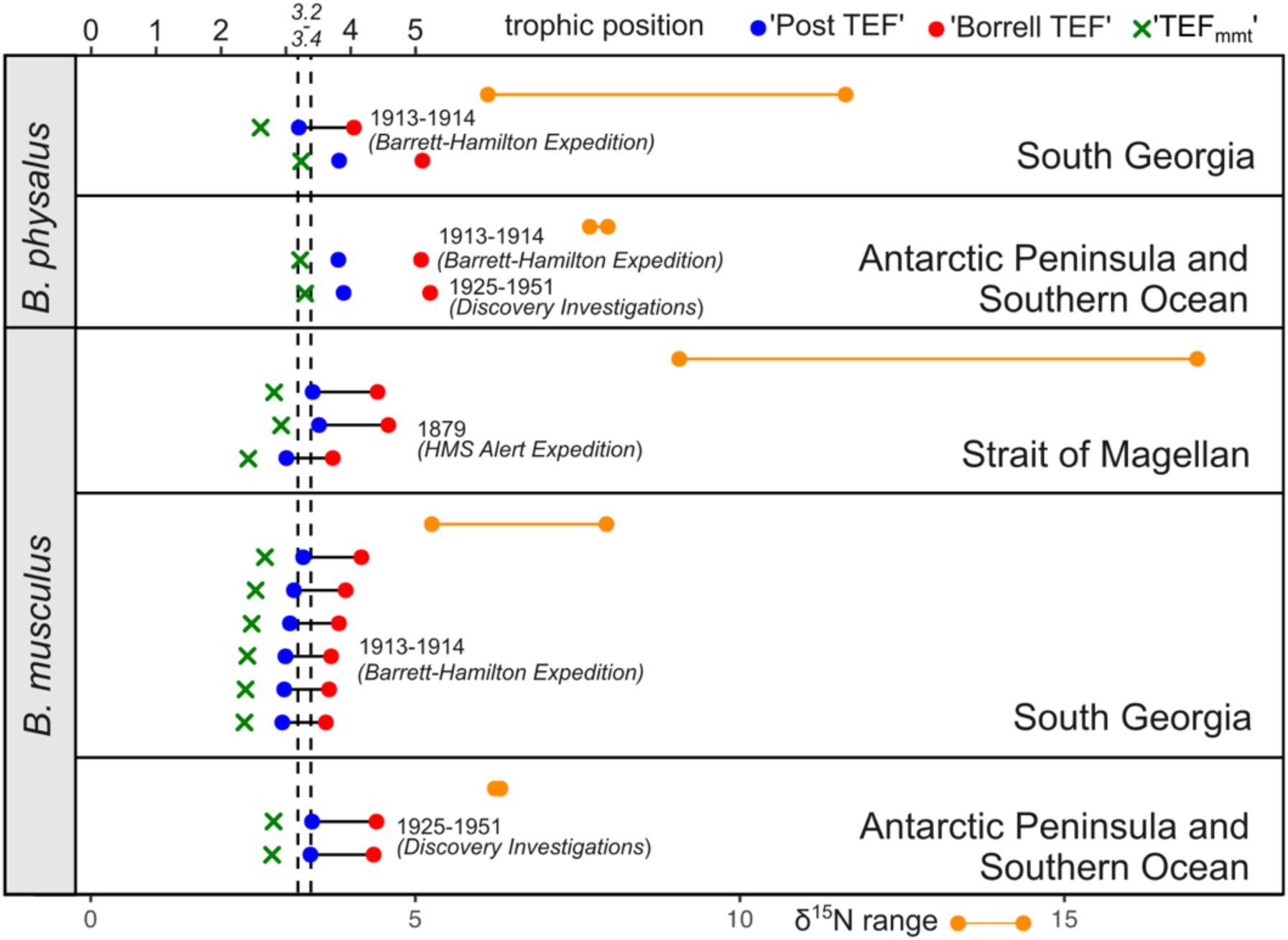
Trophic position estimates of individual baleen whale specimens and corresponding bone collagen δ^15^N ranges in each studied region. Blue and red dots, and green crosses represent three TP calculation approaches based on different TEF values (see Materials and Methods for details). Non-connected blue and red dots indicate putative geographical outliers. Line-connected orange dots mark the observed range of δ^15^N signatures in each region. Dashed lines represent expected range of trophic position for modern blue and fin whales (3.2-3.4) (51).

**Figure 4.**
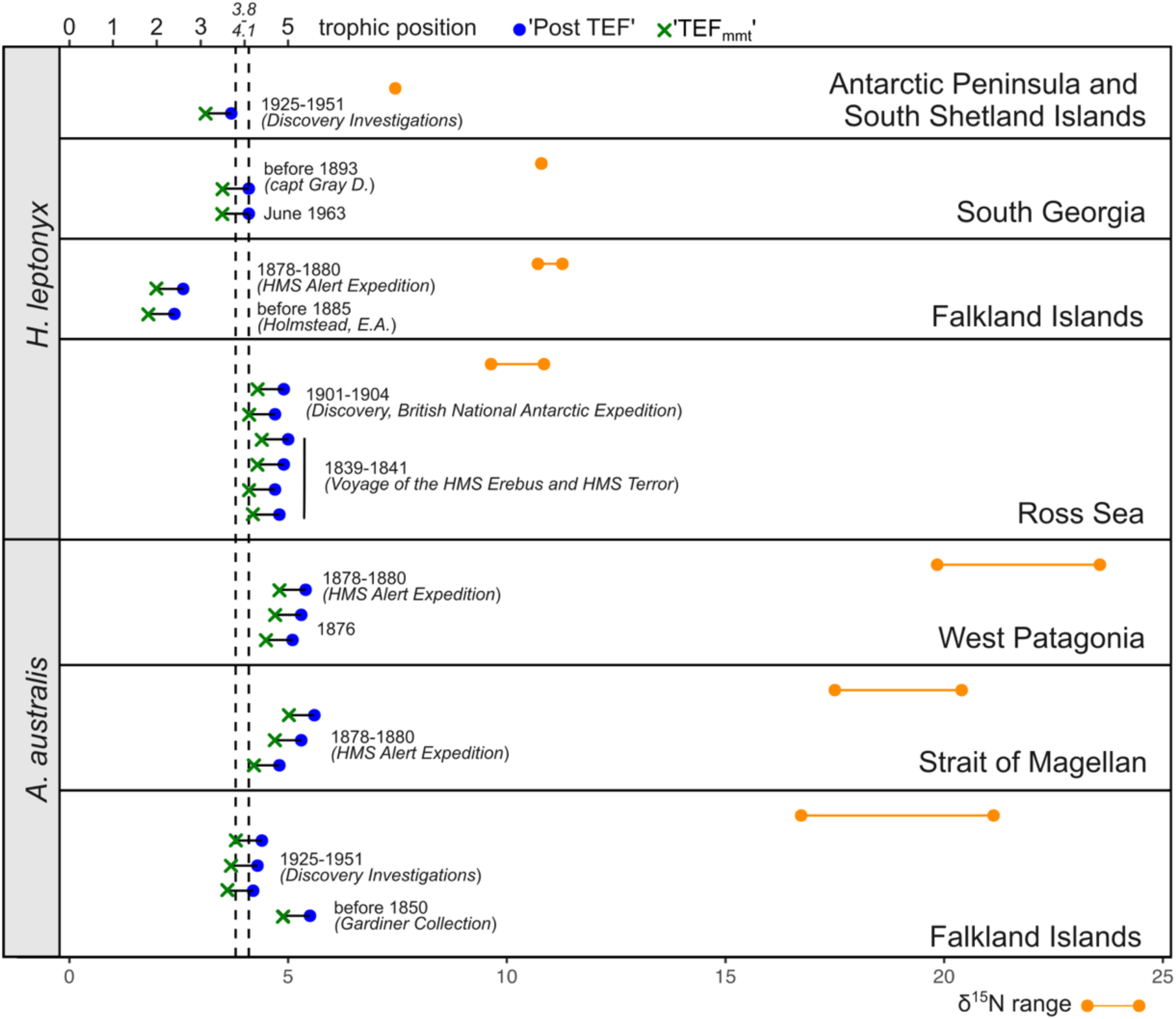
Trophic position estimates of individual leopard seal and South American fur seal specimens and corresponding bone collagen δ^15^N ranges in each studied region. Blue dots and green crosses represent three TP calculation approaches based on different TEF values (see Materials and Methods for details). Line-connected orange dots mark the observed range of δ^15^N signatures in each region. Dashed lines represent expected range of trophic position for modern seals (3.8-4.1) (51).

### Estimation of trophic positions

The calculated trophic position (TP) varies between 3.2 and 5.2 in fin whale and between 2.9 and 4.5 in blue whale; averaged TP ± SD is 3.6 ± 0.51 in blue whale and 4.3 ± 0.75 in fin whale (Figure 3), indicating several outlier values being higher or lower than the expected TP for these species. All the TP values were calculated based on the POM δ^15^N values specific to each region (See Materials and Methods for details). TP values calculated based on TEF_mmt_, i.e. corrected based on tissue-specific TEF, yield 2.4-2.9 (2.6 ± 0.21) in blue whale, and 2.6-3.3 (3.1 ± 0.32) in fin whales. Considering the high migratory abilities of the studied species and that the origin of each specimen was indicated at NHM based on the locality of landing, observed higher outlier TP values can putatively indicate more northerly/temperate foraging areas and vice versa – outliers with TP lower than expected values can indicate more southern (closer to Antarctica) foraging areas.

Comparison of TP in different time periods and regions of sampling using PERMANOVA analysis indicates significant TP differences in blue whale between the material from the 1913- 1914 (South Georgia) and 1925-1951 (Antarctic Peninsula and Southern Ocean) (PERMANOVA F=2.96, p=0.0354). Due to high within-sample variability, no differences between the 1879 samples and the subsequently collected material were revealed using PERMANOVA. No differences were also found between fin whale specimens collected in two different periods (PERMANOVA F=1.175, p=0.663).

South American fur seal TP calculated using ‘Post TEF’ varies from 4.2 to 5.6 with higher values in the Straits of Magellan and lower values in the Falkland Islands, with a mean value of 5 ± 0.53. Leopard seal TP varies between 2.4 and 5.0 (4.2 ± 0.93). The highest TP values in leopard seals are found in the Ross Sea, with lowest values in specimens from the Falklands. TP values calculated based on TEF_mmt_, vary between 3.6 and 5 (4.4±0.53) in the South American fur seals, and between 2 and 4.4 (3.6±0.93) in leopard seals.

South American fur seal specimens collected between 1850 and 1880s in the Falkland Islands, Strait of Magellan and West Patagonia do not show significant differences in TP (PERMANOVA F=0.32, p=76.84). The TP of this group of specimens collected before the 1880s significantly deviates from the TP of fur seals collected between 1925-1951 (Discovery Investigations) in the Falkland Islands (PERMANOVA F=39.85, p=0.0085). In the leopard seals, no temporal differences between the material collected in the 1830s and 1904 in the Ross Sea have been found (PERMANOVA F=0.19, p=0.6641). Comparison of all the material collected in the 19^th^ century indicates a difference in specimens from the Falklands from the other material, indicating significantly lower TP compared to other localities (PERMANOVA F=311.1, p=0.0045).

### Source prediction and foraging areas fingerprinting by MixSIAR

The Bayesian mixing modelling approach was applied to the datasets on blue whale and South American fur seal, because these species demonstrate significant changes in trophic position over the period between the 1850s and 1950s. For the blue whale specimens, mixing models were used to evaluate the contribution of various regions as key foraging areas (Figure 5). According to the MixSIAR reconstructions based on stable isotope data of five species of krill from seven geographic regions, most of the studied blue whale individuals had mixed dietary sources. The overall average dissimilarity statistics of SIMPER analysis, applied to the matrix of proportion of diet from mixing models, was 48% for the complete dataset of 11 whale specimens, with the highest contribution to the variability derived from krill from West Patagonia and South Shetland respectively (ca. 20% each).

**Figure 5.**
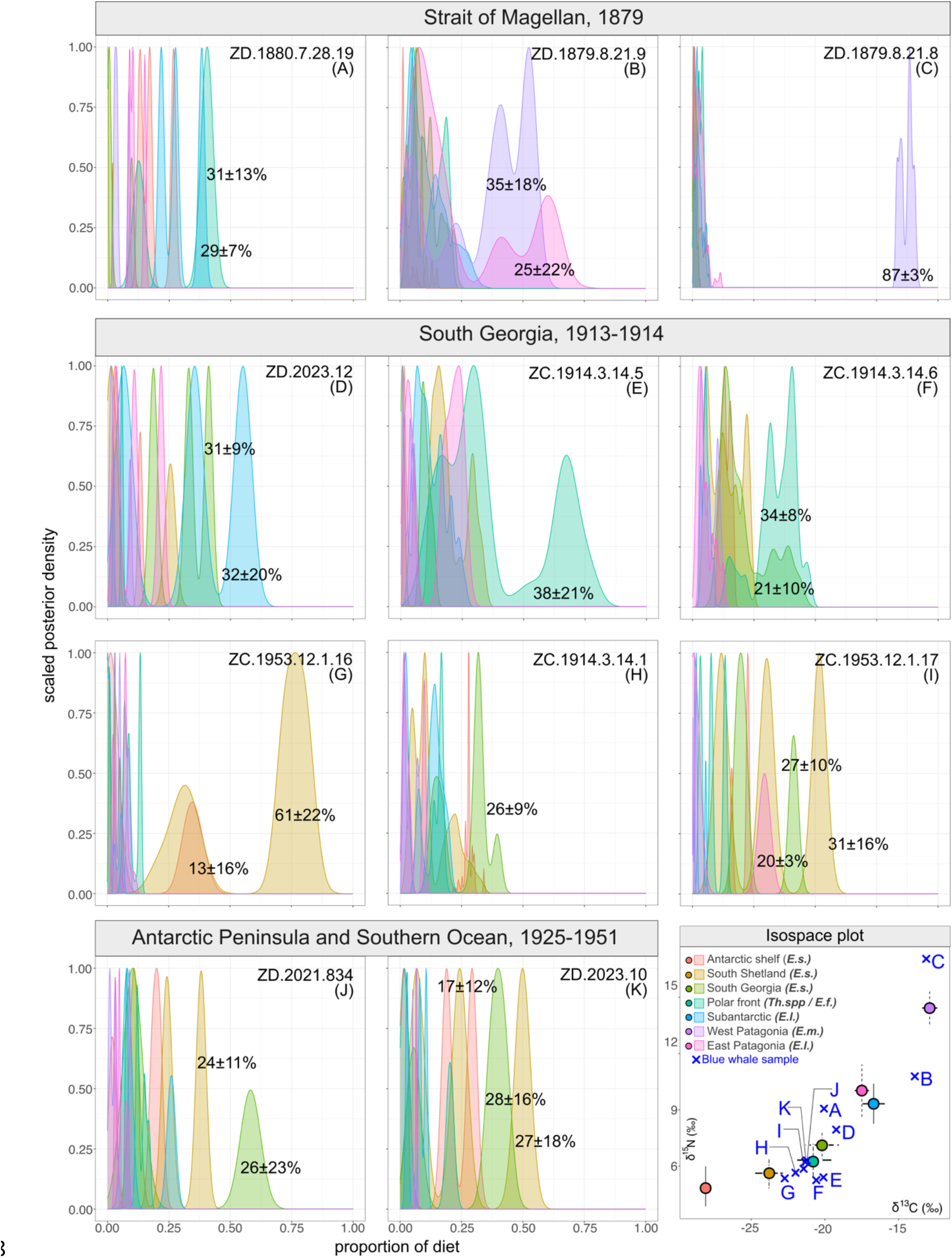
MixSIAR-estimated dietary source contributions (as Bayesian credibility intervals and posterior densities) for each sample of blue whale *B. musculus*. Abbreviations indicate potential prey items: *E.s. – Euphasia superba, Th. spp – Thysanoessa spp., E.f. – Euphasia frigida, E.l. – Euphasia lucens, E.m. – Euphasia mucronata (see Materials and methods for details)*

Among the oldest specimens in our dataset, all collected in the Strait of Magellan in the 1870s, one specimen (Figure 5C) demonstrates clear geographic affinity with the major contribution of krill from Chilean waters. For specimen ZD.1871.8.21.9 (Figure 5B), the model suggests mixed sources originating from both East and West Patagonia (i.e. South Pacific and South Atlantic). In contrast, the MixSIAR model is not able to distinguish the single major foraging area for ZD.1880.7.28.19 (Figure 5A). For the largest group of specimens in the dataset with similar geographical origin and time of collection, South Georgia (Figure 5D-I), the SIMPER overall similarity score is 41%. MixSIAR is able to predict the dominant foraging area for only one specimen of this subset (Figure 5G), the waters offshore the South Shetland Islands (61 ± 22% diet contribution). For the rest of the specimens, a relatively high contribution to the diet also contained krill from the polar front, the waters surrounding South Georgia and in the Subantarctic.

The MixSIAR model for the South American fur seals was applied to identify temporary changes in dietary sources of specimens from Falkland Islands and Strait of Magellan. According to the mixing models, the highest contribution to the diet of the specimens from both regions in the 19^th^ century was piscivorous fish species (Figure 6 A, B). On the contrary, the MixSIAR model was not able to distinguish any taxa with a major contribution to the diet for specimens dated between 1924 and 1951 from the Falkland Islands.

**Figure 6.**
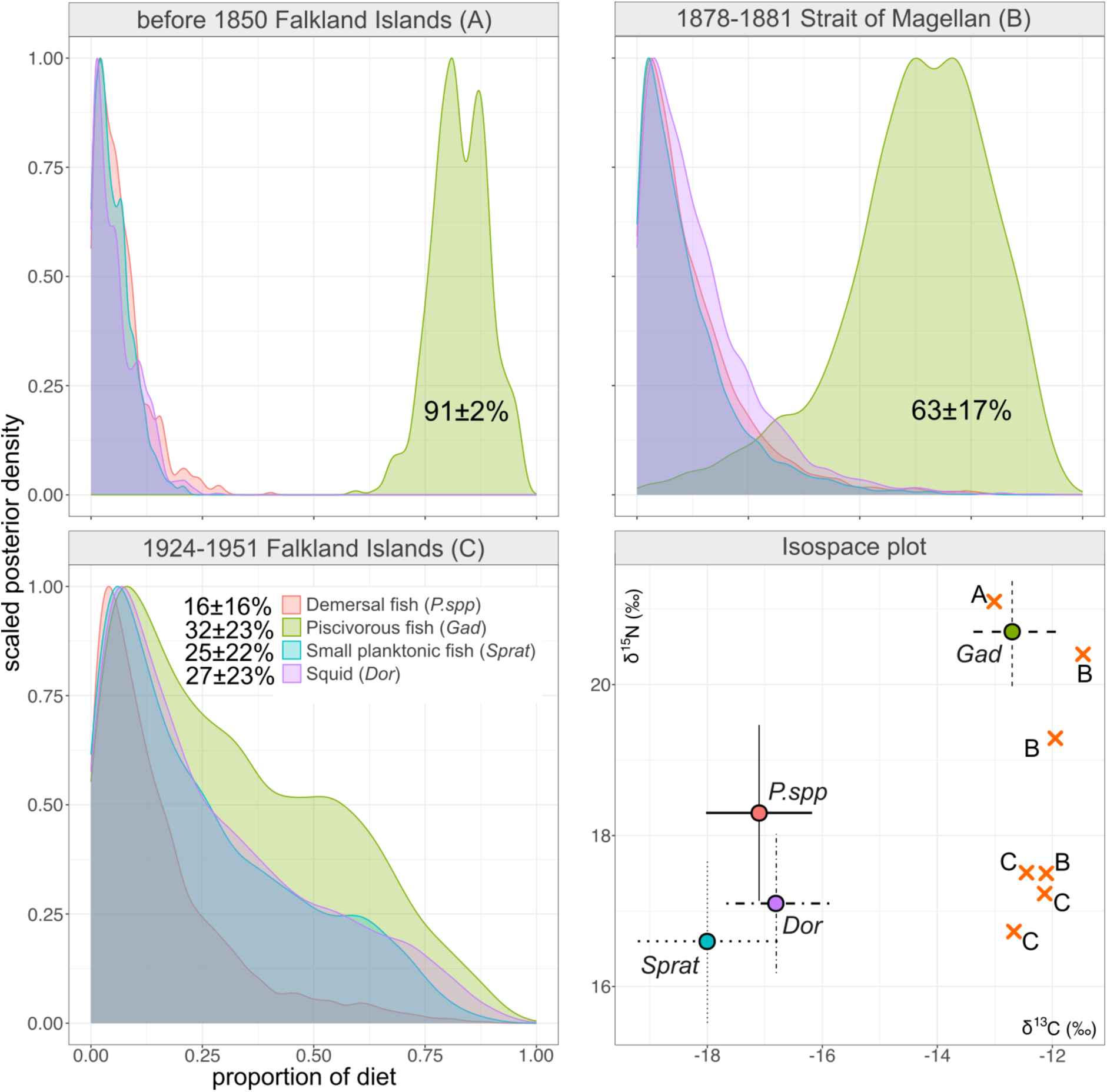
MixSIAR-estimated dietary source contributions (as Bayesian credibility intervals and posterior densities) for South American fur seal *A. australis*. *P.spp - Patagonotothen spp., Gad – Gadidae, Sprat - Sprattus fuegensis, Dor – Doryteuthis (Loligo) gahi*.

## Discussion

The number of studies examining the impacts of industrial whaling during the 20th-century is increasing (5,11,12,38,52). It is a priority to accumulate as much as possible data representing the baseline state of their populations prior to the onset of industrial-scale whaling. Despite the several uncertainties in the dataset, particularly the origin and often the unclear state of preservation of several specimens, this study provides a snapshot of stable isotope signatures of common Antarctic and Subantarctic marine mammals during the period of extensive exploration of Antarctic waters in the nineteenth century and at the turn of the 20th-century.

Museum specimens represent a unique source of information about past populations and rare species (10), but in some cases can be difficult to handle if their provenance is unclear (53). For most of the specimens used in this study, we were able to detect the geographic origin and time of collection (Table 1 Supplementary Material 2). The highest level of such uncertainty is linked to the material from the Discovery Investigations programme (1925–1951), including several specimens of blue whale, fin whale, and South American fur seal. All the samples used from the Discovery Investigations were represented as belonging to this wide time interval. Several specimens in our dataset demonstrated deviations from the commonly accepted range of C:N atomic ratio between 2.9 and 3.3 (54). Yet, since the isotopic signatures of these specimens did not deviate strongly from the other specimens from same locations and periods (Table 1, Supplementary Material 2), they were retained in our dataset; a mathematical correction for potential lipid contamination was applied to all specimens (22).

The oceanic Suess Effect is generally considered as negligible for the Southern Ocean, and thus the Suess-correction of δ^13^C often is avoided in Antarctic studies (55). A study of four species of penguins from French Southern Ocean and Antarctic territories (56) demonstrated that Suess-corrected δ^13^C values did not change over time in Antarctic species from Adelie Land, but decreased in the Subantarctic and subtropical species. However, Suess-corrected δ^13^C values of the Weddell seals from the Ross Sea decreased during the 20th century (57), indicating changes in the primary productivity of this sector of the Antarctic. Since our dataset included specimens from the Subantarctic islands, Falkland Islands and Patagonia, where the Suess Effect has been evaluated to be higher (58), we used modern isotopic signatures of POM and potential prey items from various latitudes in trophic reconstructions, the correction applied to all raw δ^13^C values and used modern datasets. In our dataset, the correction values did not alter the results based on raw δ^13^C values but were necessary for comparison of the carbon signatures among specimens from the same species collected at various locations during different periods.

We find pronounced variation in bone collagen δ^13^C and δ^15^N values across the two species of baleen whales (*B. musculus* and *B. physalus*) and in leopard seal (*H. leptonyx*). In South American fur seal (*A. australis)* we observe temporal changes in δ^15^N and no variation in δ^13^C values. The observed intraspecific variation in isotopic signatures mostly correspond to the geographic origin of specimen collection but may also potentially indicate temporal variation in foraging strategies and in the distribution of highly mobile species.

The blue whale specimens demonstrate the largest within-species variation in both carbon and nitrogen isotopes in our dataset. Among these specimens, the highest variation is observed among the 1870s data from the Magellan Strait; the isotopic signatures of these specimens is significantly different from the post-1904 dataset.

We used Bayesian mixing models as a fingerprinting tool to detect the potential foraging areas of blue whales in the South Atlantic, South Pacific and corresponding sector of the Southern Ocean. Usually, mixing models are used to evaluate the percentage contribution of potential prey items to the diet of consumers (41,59–64), and are increasingly being used to reconstruct dietary preferences of past populations from museum specimens using modern data on diet and isotopic signatures of major prey items (65–67).

For baleen whales, krill (Euphasiacea) is the main dietary source (68,69), and in the studied region taxonomic diversity and isotopic signatures of krill vary with latitude and among regions (70). The Southwest Atlantic sector of the Southern Ocean is considered as the region with highest stocks of Antarctic krill *E. superba* (71–73), High productivity and associated krill abundances are also typical of the Subantarctic islands, e.g. South Georgia (3), and across the both western and eastern Patagonian shelf (74–76). Baleen whales forage in productive areas, and various populations rely on different areas with contrasting oceanographic characteristics and prey species which can be traced using stable isotope analysis (28,77). By using stable isotope data of krill and pelagic fish from various latitudes and regions, a recent study reconstructed foraging patterns of Antarctic blue and fin whales in the Indo-Pacific sector of the Southern Ocean (55).

Mixing models suggested potential feeding ground fidelity in some of the blue whale specimens in our dataset, i.e. the accordance between the NHMUK label indicating the geographic origin of specimen and the model prediction of feeding ground. Yet, for the most specimens model predicted feeding at various locations within our study region. Two recognized subspecies of blue whales are recognised in the Southern Hemisphere and northern Indian Ocean: Antarctic (or true) blue whales (*B. musculus intermedia*) and temperate-latitude pygmy blue whale *B. musculus brevicauda*, with absolute prevalence of Antarctic blue whales to the south of 52°C (78,79). The mixing model indicate high fidelity in at least one of the 1870s whales from Strait of Magellan to Chilean sources, most likely taxonomically assigning it to the eastern South Pacific population of pygmy subspecies (80). In contrast, the whales from South Georgia and South Shetlands mostly have isotopic signatures of typical Antarctic blue whales based on their modelled dietary sources (Figure 5). Nevertheless, most of the specimens demonstrate lack of clear fidelity to one location. These results corroborate the findings of a recent study based on mark-recovery models (81). By using the Discovery Investigations and later mark data obtained between 1926 and 1963, the authors identify the high probability of blue whale movement between all sectors of Southern Ocean, indicating no mark-recovery pairs with links between Antarctic and any temperate regions (81). Our results indicate that some blue whales could potentially undertake foraging migrations between Antarctic and Patagonian waters, as mixing models predicted partial contribution of temperate and Subantarctic sources in Antarctic specimens, and vice versa, Antarctic source contribution in specimens from the Strait of Magellan. These findings do not contradict previous studies indicating that some individuals may forage at multiple locations and migrate seasonally to lower latitudes from Antarctic waters (55).

The Antarctic blue whales were hunted nearly to extinction during the 20th century, and modern population estimates are considered to be less than 1% of the pre-whaling period (82). Thus, the modern feeding patterns of blue whales in the Southern Hemisphere are likely to be not completely similar the pre-whaling period (55,81). Recent genetic studies indicate that some Antarctic blue whales demonstrate signatures of admixture with populations from temperate latitudes (52), providing a plausible basis for the interpretation of past connectivity between these populations or presence of long-distance migrants outside Antarctic waters.

The δ^15^N values from bulk stable isotope analysis cannot be compared directly among tissues derived from food webs in different areas (31,83), but the comparisons are possible through calculating trophic position based on δ^15^N values of lower trophic levels, or particulate organic matter (POM) in aquatic ecosystems (39,55,84). Our data indicate that none of the standard calculation approaches for converting bulk nitrogen isotope data to trophic position estimates can be used alone for the blue whale dataset consisting of specimens from various populations and from regions with highly contrasting productivity (Figure 3). Of the three tested, the most conventional approach (31) provided the most realistic estimates of TP compared to those previously described for the baleen whales. The applied approach of using of POM values that correspond to the region of origin of the studied whale specimens, and the observed mismatches between the expected and observed TP values show, along with the Bayesian mixing models, the mismatch between the indicated origin of specimen (e.g. landing from the whaling vessel) and potential feeding grounds in whales. Currently a more promising way of calculating the trophic position in whales and other higher marine consumers is based on data from compound-specific stable isotope analysis (CSIA) of amino acids (85). In the last decade, stable isotope analysis of specific compounds within tissues, such as individual amino acids, has become increasingly common in archaeological, trophic and animal migration studies (86,87), and when possible, comparisons of bulk and CSIA are being implemented (88).

Nevertheless, the combined use of trophic position estimations and mixing models can help explain the origin of the variable results derived from museum specimens. In our dataset, the unexpectedly high δ^15^N (17.0‰) and correspondingly high TP value (4.5) were obtained for the specimen from the Strait of Magellan, for which the mixing models predicted fidelity to the Chilean pygmy blue whale population (Figure 5C). While this δ^15^N value is very different from any of the Antarctic consumers, it is very close to values from fin whales from northern Chilean waters (17.9‰ in the winter), where nitrogen signatures of all elements of the pelagic food web are higher compared to subpolar and polar regions (89).

Our dataset includes four fin whale specimens collected between 1912 and 1951 in South Georgia, the South Shetlands and in the South Atlantic. These specimens also demonstrate pronounced variation in carbon and nitrogen isotopes, and while these results can be considered preliminary, they corroborate results from a recent baleen plate isotopic study concluding that the blue and fin whale in the Southern Ocean occupy a similar trophic level but demonstrate niche differentiation (55). One potential explanation for the observed high trophic position for several fin whale individuals is similar to those for the outlier blue whale specimens, i.e. their feeding grounds could be far from the catch locations. An additional and not mutually exclusive explanation might imply that the diet of fin whales in various parts of range include ichthyoplankton and small pelagic fish in addition to krill, and that in Patagonian waters the zooplankton consumed by whales include larvae of squat lobsters *Munida gregaria*, all occupying higher a trophic level than krill (47,55,90).

Our data indicate changes in the diet of the South American fur seal from the Falkland Islands against a background of consistent regional δ^13^C values. Estimates of trophic position reveal a significant change in the trophic preferences of fur seal in the Falkland Islands from the 19th to the 20th century. The diet of South American fur seal includes both coastal and open-sea shelf prey items (91). Currently the dominant prey items of fur seals in the Falklands, according to scat analysis, are Falkland herring *Sprattus fugensis*, and several other taxa including Patagonian squid *Doryteuthis gahi*, and several rock cod species *Patagonotothen spp.* (49). While piscivorous fish species, e.g. *Merluccius hubbsi* are rarely found in the diet of fur seals (49,91), these were included among other potential sources in our mixing model (Figure 5), primarily as an outgroup, and because recent studies based on satellite tracking data indicated long-distance foraging migrations in several colonies of South American fur seal (48). Calculation of trophic position and illustration of dietary preferences using the mixing models both indicate that South American fur seals could have had more selective feeding in the 19^th^- century, and more opportunistic feeding in the 20^th^-century, possibly due to a decline in previously preferred prey species. Seals from 1870s were presumably foraging for larger prey from higher trophic levels. As several fish taxa in Patagonian waters have recently experienced overfishing (92–94), our data could indicate that observed changes in the diet of South American fur seal represent an adaptation to changes in the pelagic ecosystem.

Leopard seal is a species capable of foraging on taxa at various trophic levels, including other seal species, penguins, fish, and krill (95–97). Huge variation in dietary preferences should affect the TP in different locations, seasons and even particular individuals. The highest TP in leopard seal was typical of specimens from the Ross Sea (4.5 ± 0.34), with samples with the lowest TP from the Falkland Islands (2.2 ± 0.36). The TP of leopard seal can be explained by contrasting foraging strategies of this species in Antarctica and the Falkland Islands. No significant temporal differences in trophic position were found in leopard seal because differences within each temporal subsets is higher that between the subsets (Figure 4). According to these data, TP of leopard seal is nearly twice lower compared to the South American fur seal. This finding is remarkable considering that fur seal pups are one of the common prey items for the leopard seal.

## CONCLUSIONS

Bulk stable isotope analyses of museum specimens in this study provide a record of Patagonian, Subantarctic and Antarctic marine ecosystems before and during the period of active exploration and exploitation of marine resources in the region at the turn of the 20th century. Here we show how this dataset can be used to address questions about the influence of industrial whaling, seal hunting and the development of fishing on the foraging of whales and seals. This study discusses the possible constraints for using this dataset in the context of conventional ecological analyses aimed at understanding the long-term changes in Antarctic marine ecosystem using several keystone species. Our results suggest that prior to and during the onset of industrial whaling in Antarctica blue whales utilized a wide variety of foraging areas with some individuals potentially undertaking foraging migrations between Antarctic and Patagonian waters. Similarly, some South Pacific blue whales could undertake foraging migrations into Subantarctic and Antarctic waters. The South American Fur seal population in the Falkland Islands could have foraged for a prey from higher trophic levels before the onset of industrial fishing in the region, and the leopard seal data demonstrate a high degree of adaptation to the local availability of prey objects. These hypotheses, based on the conventional bulk stable isotope analysis, should be further verified on the basis of larger sample sizes for each species, comparison with modern data from Antarctic and other regions within the species ranges, and using more advanced methods such as compound-specific isotope analysis that can more accurately distinguish environmental and dietary signatures.

## Acknowledgements

We thank C. Ullmann (University of Exeter) for the technical support of sampling at NHM. Phaedra Kokkini (NHMUK) provided the support during the work at NHM collection with Mattew for Tersch (BioArCh) assisted with the transfer of samples to York for the subsequent analysis. We thank David Reynolds (University of Exeter) for the useful comments during the statistical analysis of the stable isotope data and modelling.

## Ethics Statement

No approval of the research ethics committees was required to accomplish the goals of this study because of use historical specimens preserved in Natural History Museum, London (NHMUK).

## References

1. Bestley S, Ropert-Coudert Y, Bengtson Nash S, Brooks CM, Cotté C, Dewar M, et al. Marine Ecosystem Assessment for the Southern Ocean: Birds and Marine Mammals in a Changing Climate. Front Ecol Evol. 2020 Nov 4;8:566936.

2. Hammerschlag N, Williams L, Fallows M, Fallows C. Disappearance of white sharks leads to the novel emergence of an allopatric apex predator, the sevengill shark. Sci Rep. 2019 Feb 13;9(1):1908.

3. Atkinson A, Whitehouse M, Priddle J, Cripps G, Ward P, Brandon M. South Georgia, Antarctica: a productive, cold water, pelagic ecosystem. Mar Ecol Prog Ser. 2001;216:279–308.

4. Headland R. The Island of South Georgia. CUP Archive; 1984.

5. Calderan S, Black A, Branch T, Collins M, Kelly N, Leaper R, et al. South Georgia blue whales five decades after the end of whaling. Endang Species Res. 2020 Nov 19;43:359– 73.

6. Hamabe K, Matsuoka K, Kitakado T. Estimation of abundance and population dynamics of the Antarctic blue whale in the Antarctic Ocean south of 60°S, from 70°E to 170°W. Marine Mammal Science. 2023 Apr;39(2):671–87.

7. Aaris-Sørensen K, Rasmussen KL, Kinze C, Petersen KS. Late Pleistocene and Holocene whale remains (Cetacea) from Denmark and adjacent countries: Species, distribution, chronology, and trace element concentrations. Marine Mammal Science. 2010 Apr;26(2):253–81.

8. De Bruyn M, Hall BL, Chauke LF, Baroni C, Koch PL, Hoelzel AR. Rapid Response of a Marine Mammal Species to Holocene Climate and Habitat Change. Przeworski M, editor. PLoS Genet. 2009 Jul 10;5(7):e1000554.

9. English PA, Green DJ, Nocera JJ. Stable Isotopes from Museum Specimens May Provide Evidence of Long-Term Change in the Trophic Ecology of a Migratory Aerial Insectivore. Front Ecol Evol. 2018 Feb 12;6:14.

10. Smith KJ, Trueman CN, France CAM, Sparks JP, Brownlow AC, Dähne M, et al. Stable Isotope Analysis of Specimens of Opportunity Reveals Ocean-Scale Site Fidelity in an Elusive Whale Species. Front Conserv Sci. 2021 May 25;2:653766.

11. Buss DL, Hearne E, Loy RHY, Manica A, O’Connell TC, Jackson JA. Evidence of resource partitioning between fin and sei whales during the twentieth-century whaling period. Mar Biol. 2022 Nov;169(11):150.

12. Sremba AL, Martin AR, Wilson P, Cypriano-Souza AL, Buss DL, Hart T, et al. Diversity of mitochondrial DNA in 3 species of great whales before and after modern whaling. Alter E, editor. Journal of Heredity. 2023 Nov 15;114(6):587–97.

13. De Lima R, Cebuhar J, Negrete J, Ferreira A, Secchi E, Botta S. Ecosystem shifts inferred from long-term stable isotope analysis of male Antarctic fur seal Arctocephalus gazella teeth. Mar Ecol Prog Ser. 2022 Aug 25;695:203–16.

14. Szteren D, Aurioles-Gamboa D, Labrada-Martagón V, Hernández-Camacho CJ, De María M. Historical age-class diet changes in South American fur seals and sea lions in Uruguay. Mar Biol. 2018 Apr;165(4):59.

15. Deniro MJ, Epstein S. Influence of diet on the distribution of nitrogen isotopes in animals. Geochimica et Cosmochimica Acta. 1981 Mar;45(3):341–51.

16. Graham S, Mendelssohn I. Multiple levels of nitrogen applied to an oligohaline marsh identify a plant community response sequence to eutrophication. Mar Ecol Prog Ser. 2010 Nov 4;417:73–82.

17. McMahon KW, Hamady LL, Thorrold SR. A review of ecogeochemistry approaches to estimating movements of marine animals. Limnology & Oceanography. 2013 Mar;58(2):697–714.

18. Díaz A, Maturana CS, Boyero L, De Los Ríos Escalante P, Tonin AM, Correa-Araneda F. Spatial distribution of freshwater crustaceans in Antarctic and Subantarctic lakes. Sci Rep. 2019 May 28;9(1):7928.

19. Mayewski PA, Meredith MP, Summerhayes CP, Turner J, Worby A, Barrett PJ, et al. State of the Antarctic and Southern Ocean climate system. Reviews of Geophysics. 2009 Mar;47(1):2007RG000231.

20. Weiss F, Furness RW, McGill RAR, Strange IJ, Masello JF, Quillfeldt P. Trophic segregation of Falkland Islands seabirds: insights from stable isotope analysis. Polar Biol. 2009 Dec;32(12):1753–63.

21. Szpak P, Metcalfe JZ, Macdonald RA. Best practices for calibrating and reporting stable isotope measurements in archaeology. Journal of Archaeological Science: Reports. 2017 Jun;13:609–16.

22. Post DM, Layman CA, Arrington DA, Takimoto G, Quattrochi J, Montaña CG. Getting to the fat of the matter: models, methods and assumptions for dealing with lipids in stable isotope analyses. Oecologia. 2007 May;152(1):179–89.

23. Clark CT, Cape MR, Shapley MD, Mueter FJ, Finney BP, Misarti N. SuessR: Regional corrections for the effects of anthropogenic CO _2_ on δ ^13^ C data from marine organisms. Methods Ecol Evol. 2021 Aug;12(8):1508–20.

24. Eide M, Olsen A, Ninnemann US, Eldevik T. A global estimate of the full oceanic ^13^ C Suess effect since the preindustrial. Global Biogeochemical Cycles. 2017 Mar;31(3):492– 514.

25. Hobson KA, Welch HE. Determination of trophic relationships within a high Arctic marine food web using δ 13 C and δ 15 N analysis. Marine ecology progress series. 1992;9–18.

26. Lesage V, Hammill MO, Kovacs KM. Marine mammals and the community structure of the Estuary and Gulf of St Lawrence, Canada: evidence from stable isotope analysis. Marine Ecology Progress Series. 2001;210:203–21.

27. Borrell A, Abad-Oliva N, Gómez-Campos E, Giménez J, Aguilar A. Discrimination of stable isotopes in fin whale tissues and application to diet assessment in cetaceans. Rapid Comm Mass Spectrometry. 2012 Jul 30;26(14):1596–602.

28. Bury S, Peters K, Sabadel A, St John Glew K, Trueman C, Wunder M, et al. Southern Ocean humpback whale trophic ecology. I. Combining multiple stable isotope methods elucidates diet, trophic position and foraging areas. Mar Ecol Prog Ser. 2024 Apr 18;734:123–55.

29. Hobson KA, Piatt JF, Pitocchelli J. Using stable isotopes to determine seabird trophic relationships. Journal of animal ecology. 1994;786–98.

30. Minagawa M, Wada E. Stepwise enrichment of 15N along food chains: Further evidence and the relation between δ15N and animal age. Geochimica et Cosmochimica Acta. 1984 May;48(5):1135–40.

31. Post DM. Using stable isotopes to estimate trophic position: models, methods, and assumptions. Ecology. 2002;83(3):703–18.

32. Borrell A, Abad-Oliva N, Gómez-Campos E, Giménez J, Aguilar A. Discrimination of stable isotopes in fin whale tissues and application to diet assessment in cetaceans. Rapid Comm Mass Spectrometry. 2012 Jul 30;26(14):1596–602.

33. Seyboth E, Botta S, Mendes CRB, Negrete J, Dalla Rosa L, Secchi ER. Isotopic evidence of the effect of warming on the northern Antarctic Peninsula ecosystem. Deep Sea Research Part II: Topical Studies in Oceanography. 2018 Mar;149:218–28.

34. Barrios-Guzmán C, Sepúlveda M, Docmac F, Zarate P, Reyes H, Harrod C. Sample acidification has a predictable effect on isotopic ratios of particulate organic matter along the Chilean coast. Rapid Comm Mass Spectrometry. 2019 Nov 15;33(21):1652–9.

35. Bruno DO, Riccialdelli L, Acha EM, Fernández DA. Seasonal variation of autochthonous and allochthonous carbon sources for the first levels of the Beagle Channel food web. Journal of Marine Systems. 2023 Mar;239:103859.

36. Büring T, Van Der Grient J, Pierce G, Bustamante P, Scotti M, Jones JB, et al. Unveiling the wasp-waist structure of the Falkland shelf ecosystem: the role of *Doryteuthis gahi* as a keystone species and its trophic influences. J Mar Biol Ass. 2024;104:e2.

37. Espinasse B, Pakhomov E, Hunt B, Bury S. Latitudinal gradient consistency in carbon and nitrogen stable isotopes of particulate organic matter in the Southern Ocean. Mar Ecol Prog Ser. 2019 Nov 21;631:19–30.

38. Jackson J, Kennedy A, Moore M, Andriolo A, Bamford C, Calderan S, et al. Have whales returned to a historical hotspot of industrial whaling? The pattern of southern right whale Eubalaena australis recovery at South Georgia. Endang Species Res. 2020 Nov 5;43:323–39.

39. Seyboth E, Botta S, Mendes CRB, Negrete J, Dalla Rosa L, Secchi ER. Isotopic evidence of the effect of warming on the northern Antarctic Peninsula ecosystem. Deep Sea Research Part II: Topical Studies in Oceanography. 2018 Mar;149:218–28.

40. R Core Team. R: A Language and Environment for Statistical Computing [Internet]. Vienna, Austria: R Foundation for Statistical Computing; 2024. Available from: https://www.R-project.org/

41. Stock BC, Jackson AL, Ward EJ, Parnell AC, Phillips DL, Semmens BX. Analyzing mixing systems using a new generation of Bayesian tracer mixing models. PeerJ. 2018;6:e5096.

42. Stock B, Semmens B. MixSIAR GUI user manual v3. 1. Scripps Institution of Oceanography, UC San Diego, San Diego, California, USA. 2016;

43. Ciancio JE, Pascual MA, Botto F, Frere E, Iribarne O. Trophic relationships of exotic anadromous salmonids in the southern Patagonian Shelf as inferred from stable isotopes. Limnology & Oceanography. 2008 Mar;53(2):788–98.

44. Schmidt K, McClelland JW, Mente E, Montoya JP, Atkinson A, Voss M. Trophic-level interpretation based on Ã?⍰Â15N values: implications of tissue-specific fractionation and amino acid composition. Mar Ecol Prog Ser. 2004;266:43–58.

45. Riccialdelli L, Paso Viola M, Panarello H, Goodall R. Evaluating the isotopic niche of beaked whales from the southwestern South Atlantic and Southern Oceans. Mar Ecol Prog Ser. 2017 Oct 13;581:183–98.

46. Büring T, Jones JB, Pierce G, Rocha F, Bustamante P, Brault-Favrou M, et al. Trophic ecology of the squid Doryteuthis gahi in the Southwest Atlantic inferred from stable isotope analysis. Estuarine, Coastal and Shelf Science. 2023 May;284:108300.

47. Buchan SJ, Vásquez P, Olavarría C, Castro LR. Prey items of baleen whale species off the coast of Chile from fecal plume analysis. Marine Mammal Science. 2021 Jul;37(3):1116– 27.

48. Baylis A, Tierney M, Orben R, Staniland I, Brickle P. Geographic variation in the foraging behaviour of South American fur seals. Mar Ecol Prog Ser. 2018 May 28;596:233–45.

49. Baylis AMM, Arnould JPY, Staniland IJ. Diet of South American fur seals at the Falkland Islands. Marine Mammal Science. 2014 Jul;30(3):1210–9.

50. Hammer Ø, Harper D, Ryan P. Past: paleontological statistics software package for educaton and data anlysis. Palaeontologia electronica. 2001;4(1):1.

51. Pauly D, Trites A, Capuli E, Christensen V. Diet composition and trophic levels of marine mammals. ICES Journal of Marine Science. 1998 Jun;55(3):467–81.

52. Attard CRM, Sandoval-Castillo J, Lang AR, Vernazzani BG, Torres LG, Baldwin R, et al. Global conservation genomics of blue whales calls into question subspecies taxonomy and refines knowledge of population structure. Animal Conservation. 2024 Mar 15;acv.12935.

53. Keighley X, Bro-Jørgensen MH, Ahlgren H, Szpak P, Ciucani MM, Sánchez Barreiro F, et al. Predicting sample success for large-scale ancient DNA studies on marine mammals. Molecular Ecology Resources. 2021 May;21(4):1149–66.

54. Guiry EJ, Szpak P. Quality control for modern bone collagen stable carbon and nitrogen isotope measurements. Soto D, editor. Methods Ecol Evol. 2020 Sep;11(9):1049–60.

55. Smith MK, Ososky JJ, Hunt KE, Cioffi WR, Read AJ, Friedlaender AS, et al. Historical baleen plates indicate that once abundant Antarctic blue and fin whales demonstrated distinct migratory and foraging strategies. Ecology and Evolution. 2024 May;14(5):e11376.

56. Jaeger A, Cherel Y. Isotopic Investigation of Contemporary and Historic Changes in Penguin Trophic Niches and Carrying Capacity of the Southern Indian Ocean. Thrush S, editor. PLoS ONE. 2011 Feb 2;6(2):e16484.

57. Hückstädt LA, McCarthy MD, Koch PL, Costa DP. What difference does a century make? Shifts in the ecosystem structure of the Ross Sea, Antarctica, as evidenced from a sentinel species, the Weddell seal. Proc R Soc B. 2017 Aug 30;284(1861):20170927.

58. McNeil BI, Matear RJ, Tilbrook B. Does carbon 13 track anthropogenic CO _2_ in the southern ocean? Global Biogeochemical Cycles. 2001 Sep;15(3):597–613.

59. Genelt-Yanovskaya AS, Polyakova NV, Ivanov MV, Nadtochii EV, Ivanova TS, Genelt- Yanovskiy EA, et al. Tracing the Food Web of Changing Arctic Ocean: Trophic Status of Highly Abundant Fish, Gasterosteus aculeatus (L.), in the White Sea Recovered Using Stomach Content and Stable Isotope Analyses. Diversity. 2022 Nov 6;14(11):955.

60. Giménez J, Marçalo A, Ramírez F, Verborgh P, Gauffier P, Esteban R, et al. Diet of bottlenose dolphins (Tursiops truncatus) from the Gulf of Cadiz: Insights from stomach content and stable isotope analyses. Rosenfeld CS, editor. PLoS ONE. 2017 Sep 12;12(9):e0184673.

61. Guerrero AI, Pinnock A, Negrete J, Rogers TL. Complementary use of stable isotopes and fatty acids for quantitative diet estimation of sympatric predators, the Antarctic pack-ice seals. Oecologia. 2021 Nov;197(3):729–42.

62. Gül G, Demirel N. Ontogenetic shift in diet and trophic role of *Raja clavata* inferred by stable isotopes and stomach content analysis in the Sea of Marmara. Journal of Fish Biology. 2022 Sep;101(3):560–72.

63. Haug T, Biuw M, Kovacs KM, Lindblom L, Lindstrøm U, Lydersen C, et al. Trophic interactions between common minke whales (Balaenoptera acutorostrata) and their prey during summer in the northern Barents Sea. Progress in Oceanography. 2024 Jun;224:103267.

64. Mazoudier SQ, Kingsford MJ, Strickland JK, Pitt KA. Stable isotopes reveal sargassum rafts provide a trophic subsidy to juvenile pelagic fishes. Estuarine, Coastal and Shelf Science. 2023 Dec;295:108548.

65. Hiler W, Trauth SE, Wheeler B, Jimenez A, Radanovic M, Milanovich JR, et al. Stable Isotope Analysis of Ozark Hellbender (Cryptobranchus alleganiensis bishopi) Living and Preserved Museum Tissue Reveals a Shift in Their Generalist Diet Composition. Ecologies. 2021 Apr 6;2(2):187–202.

66. Torres-Poché Z, Mora MA, Boutton TW, Morrow ME. Diet sources of the endangered Attwater’s prairie-chicken in Texas: evidence from δ ^13^ C, δ ^15^ N, and Bayesian mixing models. Ecosphere. 2020 Oct;11(10):e03269.

67. Velasquez-Vacca A, Seminoff JA, Jones TT, Balazs GH, Cardona L. Isotopic ecology of Hawaiian green sea turtles (Chelonia mydas) and reliability of δ13C, δ15N, and δ34S analyses of unprocessed bone samples for dietary studies. Mar Biol. 2023 Jul;170(7):81.

68. Kawaguchi S, Atkinson A, Bahlburg D, Bernard KS, Cavan EL, Cox MJ, et al. Climate change impacts on Antarctic krill behaviour and population dynamics. Nat Rev Earth Environ. 2023 Dec 19;5(1):43–58.

69. Miller EJ, Potts JM, Cox MJ, Miller BS, Calderan S, Leaper R, et al. The characteristics of krill swarms in relation to aggregating Antarctic blue whales. Sci Rep. 2019 Nov 11;9(1):16487.

70. Schmidt K, McClelland J, Mente E, Montoya J, Atkinson A, Voss M. Trophic-level interpretation based on d15N values: implications of tissue-specific fractionation and amino acid composition. Mar Ecol Prog Ser. 2004;266:43–58.

71. Atkinson A, Siegel V, Pakhomov E, Rothery P. Long-term decline in krill stock and increase in salps within the Southern Ocean. Nature. 2004 Nov;432(7013):100–3.

72. Mascioni M, Almandoz GO, Ekern L, Pan BJ, Vernet M. Microplanktonic diatom assemblages dominated the primary production but not the biomass in an Antarctic ìord. Journal of Marine Systems. 2021 Dec;224:103624.

73. Vernet M, Martinson D, Iannuzzi R, Stammerjohn S, Kozlowski W, Sines K, et al. Primary production within the sea-ice zone west of the Antarctic Peninsula: I—Sea ice, summer mixed layer, and irradiance. Deep Sea Research Part II: Topical Studies in Oceanography. 2008 Sep;55(18–19):2068–85.

74. Antezana T. Euphausia mucronata: A keystone herbivore and prey of the Humboldt Current System. Deep Sea Research Part II: Topical Studies in Oceanography. 2010 Apr;57(7–8):652–62.

75. Franco BC, Ruiz-Etcheverry LA, Marrari M, Piola AR, Matano RP. Climate Change Impacts on the Patagonian Shelf Break Front. Geophysical Research Letters. 2022 Feb 28;49(4):e2021GL096513.

76. Nocera AC, Giménez EM, Diez MJ, Retana MV, Winkler G. Krill diel vertical migration in Southern Patagonia. Journal of Plankton Research. 2021 Jul 27;43(4):610–23.

77. Busquets-Vass G, Newsome SD, Calambokidis J, Serra-Valente G, Jacobsen JK, Aguíñiga- García S, et al. Estimating blue whale skin isotopic incorporation rates and baleen growth rates: Implications for assessing diet and movement patterns in mysticetes. Boyce MS, editor. PLoS ONE. 2017 May 31;12(5):e0177880.

78. Branch TA, Abubaker EMN, Mkango S, Butterworth DS. SEPARATING SOUTHERN BLUE WHALE SUBSPECIES BASED ON LENGTH FREQUENCIES OF SEXUALLY MATURE FEMALES. Marine Mammal Science. 2007 Oct;23(4):803–33.

79. Ichihara T. 6. The Pygmy Blue Whale, Balaenoptera musculus brevicauda, a New Subspecies from the Antarctic. In: Norris KS, editor. Whales, Dolphins, and Porpoises [Internet]. University of California Press; 1966 [cited 2025 Jan 12]. p. 79–113. Available from: https://www.degruyter.com/document/doi/10.1525/9780520321373-008/html

80. Attard CRM, Sandoval-Castillo J, Lang AR, Vernazzani BG, Torres LG, Baldwin R, et al. Global conservation genomics of blue whales calls into question subspecies taxonomy and refines knowledge of population structure. Animal Conservation. 2024 Mar 15;acv.12935.

81. Rand Z, Branch T, Jackson J. High historical movement rates of Antarctic blue whales on Southern Ocean feeding grounds estimated from Discovery mark data. Endang Species Res. 2024 Nov 14;55:109–28.

82. Shabangu FW, Stafford KM, Findlay KP, Rankin S, Ljungblad D, Tsuda Y, et al. Overview of the SOWER cruise circumpolar acoustic survey data and analyses of Antarctic blue whale calls. J Cetacean Res Manage. 2024;21–41.

83. Quillfeldt P, Masello JF. Compound-specific stable isotope analyses in Falkland Islands seabirds reveal seasonal changes in trophic positions. BMC Ecol. 2020 Dec;20(1):21.

84. Espinasse B, Pakhomov E, Hunt B, Bury S. Latitudinal gradient consistency in carbon and nitrogen stable isotopes of particulate organic matter in the Southern Ocean. Mar Ecol Prog Ser. 2019 Nov 21;631:19–30.

85. Matthews CJD, Ruiz-Cooley RI, Pomerleau C, Ferguson SH. Amino acid δ^15^ N underestimation of cetacean trophic positions highlights limited understanding of isotopic fractionation in higher marine consumers. Ecology and Evolution. 2020 Apr;10(7):3450–62.

86. Magozzi S, Thorrold SR, Houghton L, Bendall VA, Hetherington S, Mucientes G, et al. Compound-Specific Stable Isotope Analysis of Amino Acids in Pelagic Shark Vertebrae Reveals Baseline, Trophic, and Physiological Effects on Bulk Protein Isotope Records. Front Mar Sci. 2021 Sep 1;8:673016.

87. Soncin S, Talbot HM, Fernandes R, Harris A, Von Tersch M, Robson HK, et al. High- resolution dietary reconstruction of victims of the 79 CE Vesuvius eruption at Herculaneum by compound-specific isotope analysis. Sci Adv. 2021 Aug 27;7(35):eabg5791.

88. Canseco JA, Niklitschek EJ, Quezada-Romegialli C, Yarnes C, Harrod C. Comparing trophic position estimates using bulk and compound specific stable isotope analyses: applying new approaches to mackerel icefish *Champsocephalus gunnari*. PeerJ. 2024 May 17;12:e17372.

89. Andrade D, García-Cegarra AM, Docmac F, Ñacari LA, Harrod C. Multiple stable isotopes (C, N & S) provide evidence for fin whale (Balaenoptera physalus) trophic ecology and movements in the Humboldt Current System of northern Chile. Marine Environmental Research. 2023 Nov;192:106178.

90. Jory C, Lesage V, Leclerc A, Giard J, Iverson S, Bérubé M, et al. Individual and population dietary specialization decline in fin whales during a period of ecosystem shift. Sci Rep. 2021 Aug 25;11(1):17181.

91. Dassis M, Farenga M, Bastida R, Rodríguez D. At-sea behavior of South American fur seals: Influence of coastal hydrographic conditions and physiological implication. Mammalian Biology. 2012 Jan;77(1):47–52.

92. Bingham M. The decline of Falkland Islands penguins in the presence of a commercial fishing industry. Revista chilena de historia natural. 2002;75(4):805–18.

93. Heredia-Azuaje H, Niklitschek E, Sepúlveda M, Harrod C, Guerrero A, Peña G, et al. Salmon farming, overfishing and southern sea lion: Not so opportunistic responses of a top predator to human perturbations in the Patagonian Fjords. Estuarine, Coastal and Shelf Science. 2024 Apr;299:108669.

94. Tschopp A, Herrera VP, García NA, Crespo EA, Coscarella MA. Temporal changes in the diet composition of Shorttail Yellownose skate, when exposed to overfishing conditions in northern and central Patagonia, Argentina. Hydrobiologia. 2024 Jun;851(11):2695– 710.

95. Boveng PL, Hiruki LM, Schwartz MK, Bengtson JL. POPULATION GROWTH OF ANTARCTIC FUR SEALS: LIMITATION BY A TOP PREDATOR, THE LEOPARD SEAL? Ecology. 1998 Dec;79(8):2863–77.

96. Krause DJ, Goebel ME, Kurle CM. Leopard seal diets in a rapidly warming polar region vary by year, season, sex, and body size. BMC Ecol. 2020 Dec;20(1):32.

97. Sperou ES, Crocker DE, Borras-Chavez R, Costa DP, Goebel ME, Kanatous SB, et al. Large and in charge: cortisol levels vary with sex, diet, and body mass in an Antarctic predator, the leopard seal. Front Mar Sci. 2023 Jun 23;10:1179236.

